# A mesh microelectrode array for non-invasive electrophysiology within neural organoids

**DOI:** 10.1101/2020.09.02.279125

**Authors:** Matthew McDonald, David Sebinger, Lisa Brauns, Laura Gonzalez-Cano, Yotam Menuchin-Lasowski, Michael Mierzejewski, Olympia-Ekaterini Psathaki, Angelika Stumpf, Jenny Wickham, Thomas Rauen, Hans Schöler, Peter D. Jones

## Abstract

Organoids are emerging *in vitro* models of human physiology. Neural models require the evaluation of functional activity of single cells and networks, which is best measured by microelectrode arrays. The characteristics of organoids clash with existing *in vitro* or *in vivo* microelectrode arrays. With inspiration from implantable mesh electronics and growth of organoids on polymer scaffolds, we fabricated suspended hammock-like mesh microelectrode arrays for neural organoids. We have demonstrated the growth of organoids enveloping these meshes and the culture of organoids on meshes for up to one year. Furthermore, we present proof-of-principle recordings of spontaneous electrical activity across the volume of an organoid. Our concept enables a new class of microelectrode arrays for *in vitro* models of three-dimensional electrically active tissue.

## 1 Introduction

Neural organoids show promise as physiologically relevant *in vitro* models of the human brain.^1^ Electrophysiology is critical to demonstrate functionality of translational models of the central nervous system.^2^ Electrical activity has been observed in organoids by adapting existing tools (Figure 1B–F), however, there remains an unmet need for more appropriate tools and methods specific for measuring the electrophysiological properties of organoids.^3^

**Figure 1:**
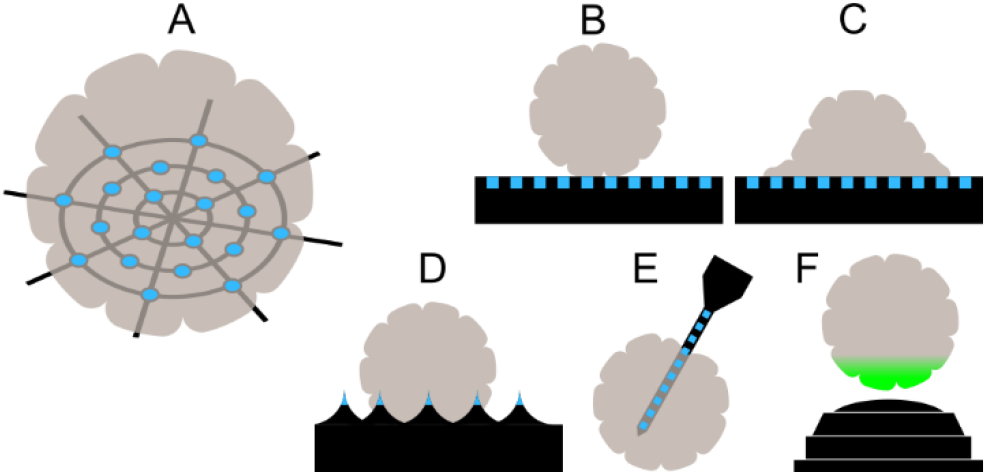
Technical possibilities for recording electrical activity of neural organoids. A: Mesh MEA (this work). B: Acute use of a planar MEA. C: Culture on a planar MEA. D: Planar MEA with protruding electrodes. E: Probe implantation. F: Calcium or voltage imaging.

What is needed for measuring organoid electrophysiology? Scientists may have different demands: Can we extract relevant information using only a handful of electrodes?^4^ Can we measure the activity of all neurons in an organoid,^5^ and would this be useful? How does the activity of organoids relate to human physiology?^1^ Should we consider new ethical concerns?^6^ In fact, current knowledge of the functional electrical activity within organoids does not allow these questions to be addressed and thus we suggest a new technical approach to tackle them.

Neural organoids are on a spectrum between conventional cell culture and a living brain. Research tools are available for both ends of this spectrum, yet are not ideal for organoids. In conventional cell culture, cells can adhere onto planar microelectrode arrays (MEAs). Typical *in vitro* models relate action potentials and bursting activity to cellular or synaptic effects of applied compounds. In contrast, animal models attempt to resolve the activity of many single neurons over an extended period of time, and to relate it to parameters such as behavior.

In either case, microelectrodes may be sparsely distributed to efficiently capture independent activity or may be densely packed to capture all activity at a high resolution. In the simplest *in vitro* models, network activity can be captured by only a few electrodes, yet many independent samples are needed to measure small effects.^4^ In the brain, we expect complex network activity; an ultimate vision in neuroscience is to record all cells at millisecond resolution *in vivo*.^5,7^ We assume organoids could fall anywhere between these two extremes. While existing MEAs are appropriate for simple organoids, more advanced MEAs are needed for capturing the complex activity that is expected in highly structured organoids.

Capturing the activity of single cells requires close proximity to electrodes – and tracking cells over time requires a stable structure. Therefore, an alternative to conventional methods of growing freely floating organoids is needed. Organoids can be grown on 2D-MEAs for several months^8^ and such MEAs can detect activity to a depth of 100 μm.^9^ While round organoids may contact few electrodes on a planar MEA (Figure 1B), organoids can flatten as neurons migrate and spread on the MEA surface (Figure 1C). The ability to resolve internal activity similar to electroencephalography (EEG) cannot be assumed in organoids, as EEG measures coordinated activity in structured cell layers.^10^ Deeper activity can be recorded by MEAs with protruding microelectrodes (Figure 1D) and slicing of organoids^11^ or implantable neural probes (Figure 1E). In addition to the technical challenge of implantation, tissue damage limits recordings to a single endpoint.^12^ Artificial devices that block diffusion of oxygen will increase hypoxic stress in the cells nearest to the device (Figure 1B–D). Beyond such electrical methods, optical methods can record superficial activity (Figure 1F) but with lower sensitivity.^13^

To move beyond these limited approaches, we were inspired by the growth of organoids on artificial structures^14^ and demonstrations of implantable mesh electronics with cellular or subcellular structure sizes.^15,16^ We believe that the combination of these approaches should enable guided growth of neural organoids with non-invasive electrophysiology (Figure 1A).

A similar device has been demonstrated in acute tissue experiments and suggested for organoids but has only 16 or 32 high-impedance gold microelectrodes and is incompatible with long-term cultures.^17^ Superficial electrophysiology of cardiac spheroids has been demonstrated with a flexible self-rolling device, but the non-mesh structure restricts diffusion and organoid growth.^18^

In this work, we present a mesh microelectrode array specifically designed for physically suspended growth and long-term electrophysiology of neural organoids (Figure 1A). With a hammock-like structure, the device suspends organoids far from solid surfaces to enable their unimpeded growth. Our successful extracellular recording of spontaneous action potentials supports the use of such devices for non-invasive electrophysiology of activity within and throughout neural organoids. By simultaneously measuring the electrical activity at many sites within neural organoids, these mesh MEAs will enable advanced studies of the network activity in highly structured organoids.

## 2 Methods

### 2.1 Design

In designing this organ-on-a-chip device, we prioritized compatibility with organoid culture methods, minimization of technical complexity and material choices for long-term device stability. We adopted methods previously used for fabrication of perforated MEAs.^19^ Microelectrodes (30 μm diameter) were sparsely distributed across a suspended region with a diameter of 2 mm. Minimum dimensions of 4 μm (lines and spaces) ensured a high fabrication yield. We focused on a design compatible with 256-channel amplifiers from Multi Channel Systems MCS GmbH (Reutlingen) with minimal complexity of electrical packaging. Electrical connections are made by gold pins of the amplifier to peripheral contact pads.

### 2.2 Fabrication

Our devices consist of flexible microfabricated meshes assembled into glass-bottomed wells (Figure 2). The meshes were produced on carrier wafers using cleanroom microfabrication by NMI TT GmbH (Reutlingen, Germany).^19^ Electrical paths (Ti/Au/Ti, 400 nm) were insulated above and below by 6 μm of polyimide (DuPont PI-2611). Metals were deposited by sputter coating and structured by plasma etching. Polyimide was structured by plasma etching against a silicon nitride hard mask. Microelectrodes (30 μm diameter) of titanium nitride (TiN, 500 nm) were structured by sputter deposition and lift-off using a photoresist mask. After release from carrier substrates, we assembled the meshes into wells comprising multiple machined polymer and glass components using the adhesive EPOTEK 301-2SL (Figure 2E). Glass components were pretreated with aminopropyltriethoxysilane before gluing.

**Figure 2:**
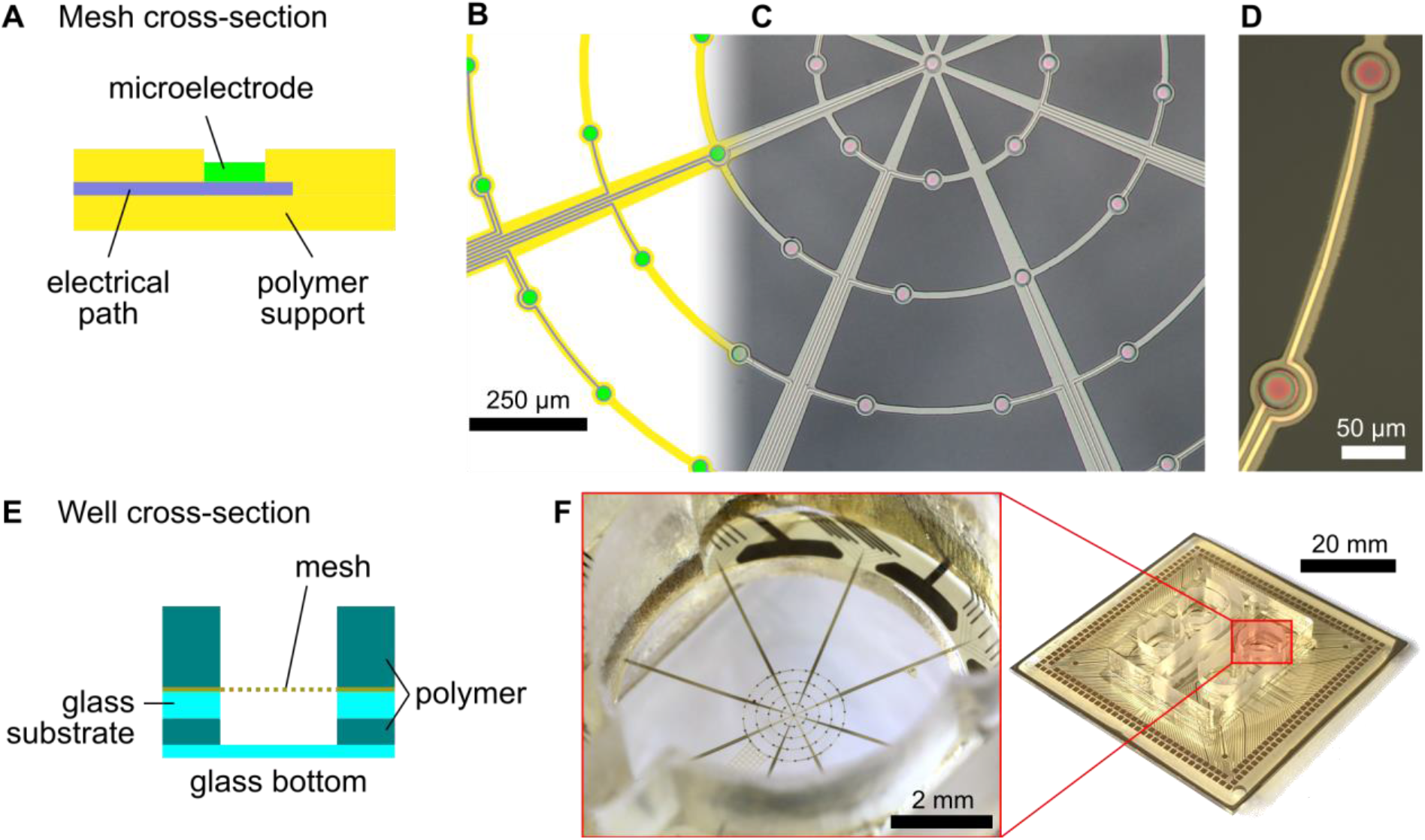
Mesh MEA device. A: Schematic cross-section of the polymer filaments. B, C: Drawing and photo of the suspended mesh. Regions between polymer filaments are open. D: Microelectrodes on a single filament. E: Schematic cross-section of a well. The well has an inner diameter of 7 mm. The glass bottom is 0.17 mm thick. The lower polymer piece and glass substrate are each 1 mm thick. The mesh is 12 μm thick. The top polymer piece is 5 mm thick. F: Photos of the whole device and a single well with a suspended hammock-like mesh.

### 2.3 Cell and organoid culture

In this paper, we present results of two types of neural organoids generated from human induced pluripotent stem (iPS) cells. Cells were verified pluripotent and contamination-free, confirmed negative for mycoplasma, and pluripotent stem cells were maintained feeder-free. Our methods for culturing organoids on the mesh MEA were exploratory and should not be considered as best practices.

Cerebral organoids (Figure 3) were generated following the protocol published by Lancaster et al.^20^ Human iPS cells were obtained in house via OSKM reprogramming.^21^ Briefly, at day 0, 7000 cells from single cell suspensions were plated in 96-well U-bottom low attachment plates to generate embryoid bodies. After 6 days in vitro, media composition was changed to induce the formation of anterior neuroectoderm. At day 10 the neuroectodermal aggregates were embedded in Matrigel to provide them with a scaffold that would allow for further differentiation, maturation and growth.

**Figure 3:**
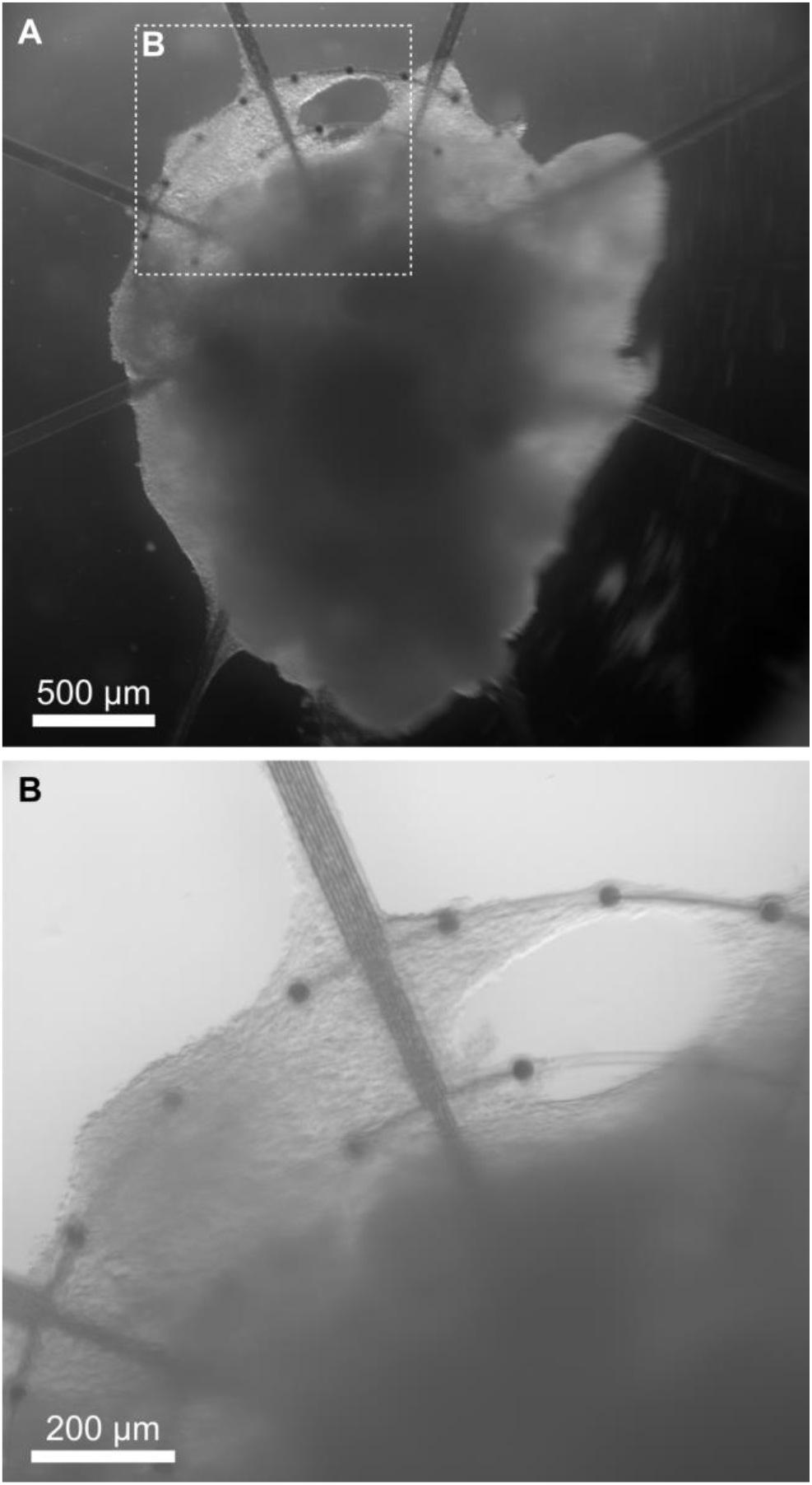
An organoid growing on a mesh MEA. This 40-day-old neural organoid showed outgrowth of cells along the polymer filaments after ten days on the mesh. Thin tissue reveals embedded microelectrodes (black circles in B). Optical images were taken from above.

For a second type of neural organoid (Figure 4, Figure 5), human iPS cells were obtained from Thermo Fisher Scientific (A18945). On day 0 of organoid culture, iPS cells (passage < 50, approximately two thirds confluence) were dissociated from plates by EDTA (Sigma) treatment and plated in ultra-low-binding plates (Corning) containing mTeSR™1 medium (STEMCELL Technologies) with blebbistatin (Calbiochem) added. Neural induction was induced by gradually exchanging mTeSR™1 medium with neural induction medium [DMEM/F12 (Gibco), N-2 supplement (Gibco), Glutamax (Gibco), MEM-NEAA (Gibco) and heparin (Sigma)] from day 1 on. At day 9, tissues were embedded into Matrigel (Corning) and thereafter grown in differentiation media [DMEM/F12 and Neurobasal (Gibco) containing N-2 supplement, B27 supplement (Gibco), 2-mercaptoethanol (Sigma), insulin (Sigma), Glutamax, penicillin-streptomycin (Gibco) and MEM-NEAA].

**Figure 4:**
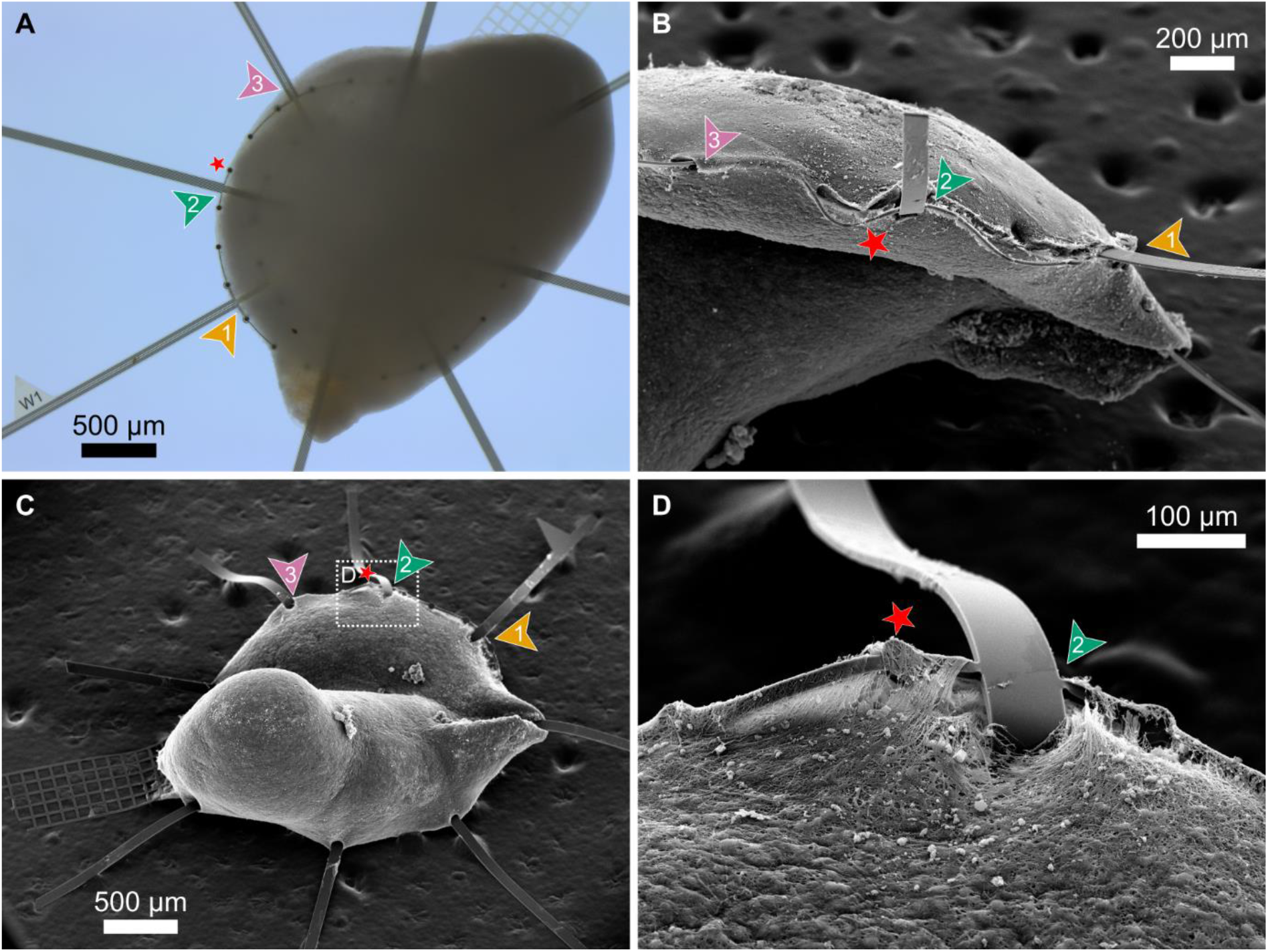
Organoids envelope the mesh structure. A: An optical image taken from above. B–D: SEM images taken from the side and below reveal how this organoid grew above and below the mesh. Arrows indicate three filaments to assist with orientation. A star indicates the same microelectrode in all images. B shows the microelectrode from the side, while C and D are from below. In D, the polyimide surface below the microelectrode is visible. The majority of microelectrodes are concealed within the bulk of the organoid. This organoid grew on the mesh for one year.

**Figure 5:**
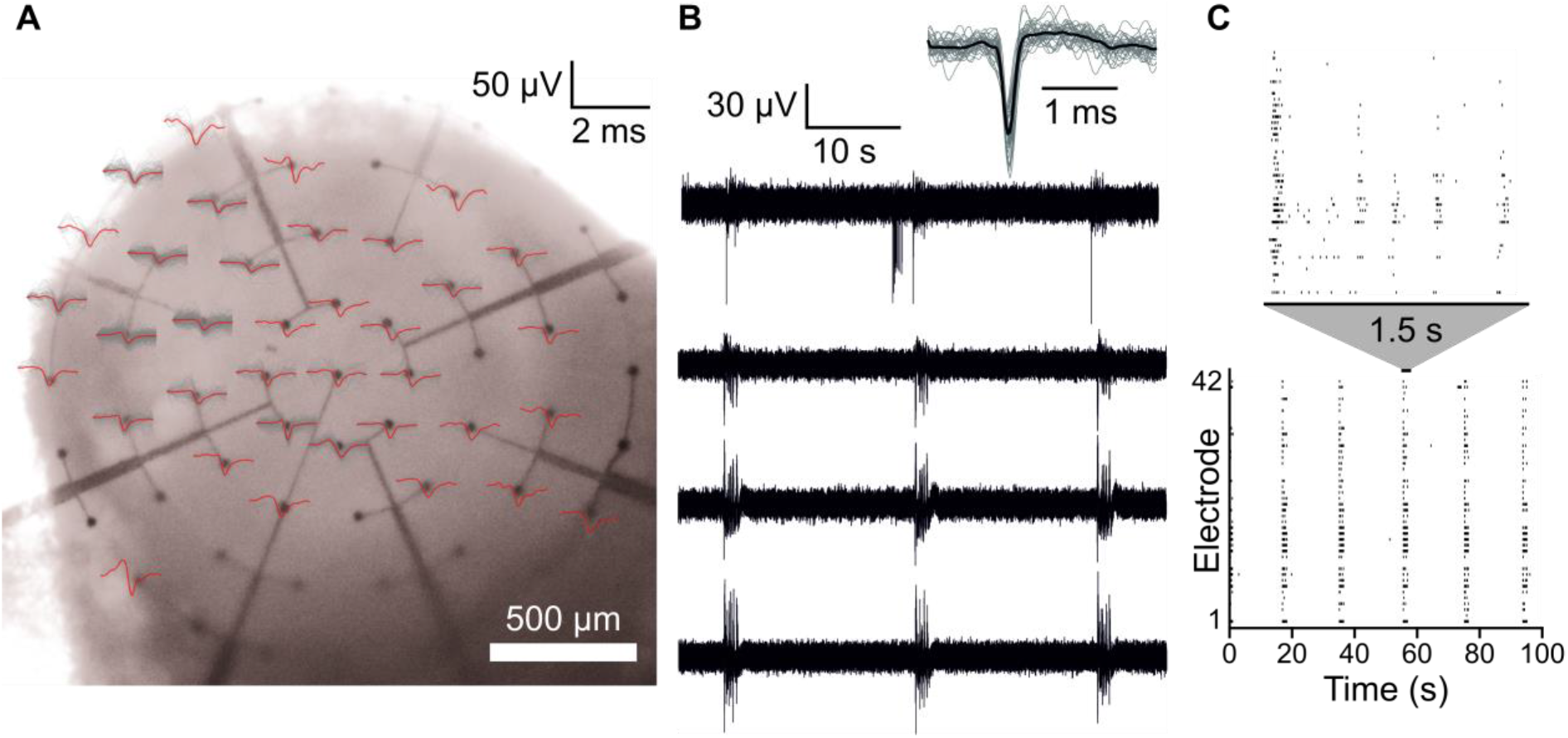
Electrical activity in a neural organoid, shown with detected waveforms overlaid at electrode positions (A; raw spikes in grey and average in red), example traces from four electrodes together with an average waveform of the action potentials from the first trace (B) and a spike raster plot (C).

We transferred individual organoids onto mesh MEAs by pipetting. For cerebral organoids, low media volumes for the first 5 days on the mesh kept the organoids at the air–liquid interface to facilitate enveloping of the mesh. Later, full media changes covering the entire organoid were performed every other day. Before placing our second type of organoids, MEAs were coated with poly-D-lysine to promote cell adhesion: MEAs were treated with air plasma, incubated with 1 mg/ml poly-D-lysine for 1 h, then rinsed three times with saline. Media was changed every other day and maintained to keep organoids at the air–liquid interface.

### 2.4 Electrophysiology

Spontaneous electrical activity was recorded by a multichannel amplifier (USB-MEA256 from Multi Channel Systems MCS GmbH) at a sampling rate of 40 kHz. During offline analysis recordings were high-pass filtered (200 Hz second-order Butterworth) and action potentials were identified by threshold detection.

### 2.5 Electron microscopy

For scanning electron microscopy (SEM) organoids were fixed for 2 h at room temperature in 2.5 % glutaraldehyde (Sigma, Germany) in 0.1 M cacodylate buffer at pH 7.4 (Sciences Services, Germany), subsequently washed twice in 0.1 M cacodylate buffer at pH 7.4 and dehydrated stepwise in a graded ethanol series. Specimens were critical point dried in 100 % ethanol (CPD 300, Leica, Austria) and afterwards glued onto conductive carbon pads (Leit-Tabs, Plano, Germany) mounted on aluminum stubs (Plano, Germany) and were sputter-coated with a 15 nm gold layer (ACD600, Leica, Austria). SEM images were acquired with a JEOL JSM-IT200 SEM (JEOL, Japan) at 15 kV with a secondary electron detector.

## 3 Results and discussion

### 3.1 Device

The devices (Figure 2) contain four wells with diameters of 7 mm. Each well can support a single organoid on a central spider-web-like mesh with a diameter of 2 mm and 61 microelectrodes. The 30-μm-diameter TiN microelectrodes enable low noise recordings^22^ (2-3 μV_rms_) and are suitable for electrical stimulation.^23^ Each mesh has four concentric ring filaments (radii of 0.25, 0.5, 0.75 and 1 mm) containing 8, 12, 16 and 24 electrodes, and one electrode at the center of the mesh. The rings are suspended 2 mm above the glass bottom. Eight radial filaments provide structural support and electrical connections to the electrodes. The filaments have a thickness of 12 μm and width of 20 μm. The width of the radial filaments increases to 80 μm to accommodate all electrical connections. Outside of the wells, the electrical paths extend to gold pads at the device perimeter, which are contacted directly by the amplifier headstage.

We designed this device to allow unrestricted growth in all directions, in contrast to the perturbed growth and restricted oxygen supply of organoids on planar MEAs. In the suspended plane of the mesh, 85 % of the central 2 mm circle is open space. With its thickness of 12 μm, the artificial structure would occupy only 0.1 % of the total volume of a spherical 2-mm-diameter organoid. The impact on the development of such an organoid should be greatly reduced in comparison to organoids grown on planar MEAs. Suspending the organoid prevents its adhesion on solid surfaces, thereby reducing the need for mechanical shaking. Nutrients and oxygen can be delivered from all sides, although perfusion could improve delivery. No flow was used in the presented results.

The sparse layout of electrodes maximizes the volume from which action potentials may be detected. Each electrode may detect action potentials from soma as distant as 100 μm.^5^ With inter-electrode distances on the order of 200 μm, the majority of action potentials generated near the plane of the mesh should be detected. A disadvantage of a sparse design is that activity of any cell should be recorded by at most one electrode, which prevents spatial triangulation and can limit spike sorting.

We designed our mesh to support microelectrodes in a fixed arrangement for simple placement of organoids and electrode localization even when embedded in tissue. However, the materials used to produce the mesh would allow flexibility or stretching if modified to include serpentine structures.^24^ Our microelectrodes are connected by unbroken thin film metal paths rather than complicated electrical packaging.^25,17,24^ The microfabricated structures of our device are similar to the flexible devices reported by Soscia et al., although their device is intended for 3D cultures of cells suspended in hydrogel.^26^ In comparison to the use of rigid devices for recording in 3D cell cultures,^11,12^ devices such as ours or those of Soscia et al. which integrate electrodes on flexible filaments may allow for a better long-term culture of organoids.

We have observed no problems of leakage or technical instability, even after culture of organoids for up to one year.

### 3.2 Organoids

After their placement on a mesh MEA, organoids enveloped the filaments and continued to grow. Figure 3 shows a cerebral organoid, which had grown on the mesh for ten days. The decentered placement of this example – although unintentional – revealed how an organoid can grow on filaments of a mesh MEA. Thin regions of tissue reveal the embedded microelectrodes, while thicker regions hide the filaments and microelectrodes. Figure 4 shows how an organoid (of our second type) placed on the mesh grew above and below the mesh. This organoid, whose electrical activity is shown in Figure 5, grew on the mesh for one year.

A limitation of our design is the large open spaces peripheral to the central electrode field. Adhesion of organoids after placing them on meshes is not immediate, but requires the organoids to be returned to an incubator. Even careful handling caused some organoids to fall off their meshes, depending on their shape, type and size. We are filling these regions with mesh in newer devices, although a user-friendly solution such as well inserts may simplify placement of organoids.^27^

How an organoid interacts with the mesh will depend on its physical properties, shape and mechanical characteristics and also its biological maturity and behavior. Possible interactions with the mesh include adhesion, migration of cells along the artificial structures, and engulfment by rearrangement or proliferation of cells. Best practices may depend on the organoid type and the purpose of the model. We envision that growth on the mesh from earlier states (embryoid bodies or stem cells) may be possible. However, this will require additional adaptation of biological protocols.

### 3.3 Electrophysiology

Three neural organoids derived by our second method were placed on three meshes at an age of 197 days. Recordings of spontaneous activity were performed 35 days later (day 232). We observed spontaneous activity on most microelectrodes across a distance of 2 mm in one organoid (Figure 5A) and action potentials could be detected using threshold crossings (Figure 5B). Synchronized bursting activity similar to ref. 8, recurring continuously over the whole recording session, was measured on most electrodes (Figure 5C). This activity indicates the formation of a synaptically connected neuronal network. We measured activity in the same organoid one month and eight months later. After one year on the device, this organoid was imaged by SEM (Figure 4). Signal amplitudes decrease with increasing distance between electrodes and active cells,^9^ and our measured amplitudes suggest that active cells are not directly adjacent to the microelectrodes. Time-dependent studies to compare cell types and morphologies with electrophysiology will be possible by histology of organoids containing embedded mesh microelectrode arrays.^16^

We present these electrophysiological results as a demonstration of the device’s functionality, but caution against generalizing these preliminary results to other organoids. Interestingly, the two sibling organoids did not develop spontaneous activity. Likewise, we also measured an absence of electrical activity for cerebral organoids and comment on possible causes below.

It cannot be assumed that neural organoids will be electrically active per se. The emergence of functional activity may be limited by variations in differentiation or cultivation protocols. Activity could be suppressed by culture medium composition, or changes in gas composition or culture temperature during recordings.^28^ Devices which restrict oxygen diffusion or cause damage to organoids (Figure 1) may suppress neuronal activity. Recordings of spontaneous action potentials depend on sufficient levels of intrinsic activity as well as close proximity between spiking neurons and microelectrodes. Furthermore, activity may emerge and again disappear as an organoid and its constituent cells develop. Troubleshooting an absence of electrical activity can therefore be challenging – especially considering the long culture period required for organoid formation and maturation.

## 4 Conclusions

We have constructed a microelectrode array device specifically for long-term cultivation and non-invasive electrophysiology within neural organoids. With such devices, we hope to contribute to a better understanding of electrophysiological activity in organoids. Enabling functional read-out will help to make neural organoids useful models for human neurodevelopment and disease.^1,2^

Full benefits of organoid electrophysiology may be obtained by considering both the device design and organoid culture techniques. Methods which place organoid precursors such as embryoid bodies or stem cells may require modified devices. Assembly of organoids of various brain regions or other tissues may be enabled by the development of specific designs. Future implementations may balance the numbers of wells and microelectrodes for targeted applications. Further technical advancements could include stretchable meshes, mechanical or chemical sensors, or three-dimensional meshes.

## 6 Acknowledgements

We thank Andrea Corna, Ricarda Stock, Dr. Hansjürgen Volkmer and Dr. Günther Zeck for helpful discussions. We thank Dr. Clemens Boucsein and Dr. Frank Hofmann from Multi Channel Systems MCS GmbH for providing a 256-channel amplifier. We acknowledge funding from the Max Planck Society’s White Paper Project “Alternatives to Animal Testing,” the Federal Ministry of Education and Research (BMBF) for support under grant 031L0059A (DREPHOS), and the German Research Foundation (DFG) under SFB 944 Z-Project.

